# Comparative analysis between single-cell RNA-seq and single-molecule RNA FISH indicates that the pyrimidine nucleobase idoxuridine (IdU) globally amplifies transcriptional noise

**DOI:** 10.1101/2023.03.14.532632

**Authors:** Giuliana P. Calia, Xinyue Chen, Binyamin Zuckerman, Leor S. Weinberger

**Affiliations:** Gladstone|UCSF Center for Cell Circuitry, University of California, San Francisco, CA 94158; Department of Biochemistry and Biophysics, University of California, San Francisco, CA 94158; Department of Pharmaceutical Chemistry, University of California, San Francisco, CA 94158

**Keywords:** noise enhancer molecule, transcriptional noise, single-cell RNA sequencing

## Abstract

Stochastic fluctuations (noise) in transcription generate substantial cell-to-cell variability, but the physiological roles of noise have remained difficult to determine in the absence of generalized noise-modulation approaches. Previous single-cell RNA-sequencing (scRNA-seq) suggested that the pyrimidine-base analog (5’-iodo-2’-deoxyuridine, IdU) could generally amplify noise without substantially altering mean-expression levels but scRNA-seq technical drawbacks potentially obscured the *penetrance* of IdU-induced transcriptional noise amplification. Here we quantify global-vs.-partial penetrance of IdU-induced noise amplification by assessing scRNA-seq data using numerous normalization algorithms and directly quantifying noise using single-molecule RNA FISH (smFISH) for a panel of genes from across the transcriptome. Alternate scRNA-seq analyses indicate IdU-induced noise amplification for ~90% of genes, and smFISH data verified noise amplification for ~90% of tested genes. Collectively, this analysis indicates which scRNA-seq algorithms are appropriate for quantifying noise and argues that IdU is a globally penetrant noise-enhancer molecule that could enable investigations of the physiological impacts of transcriptional noise.

**Graphical Abstract:** 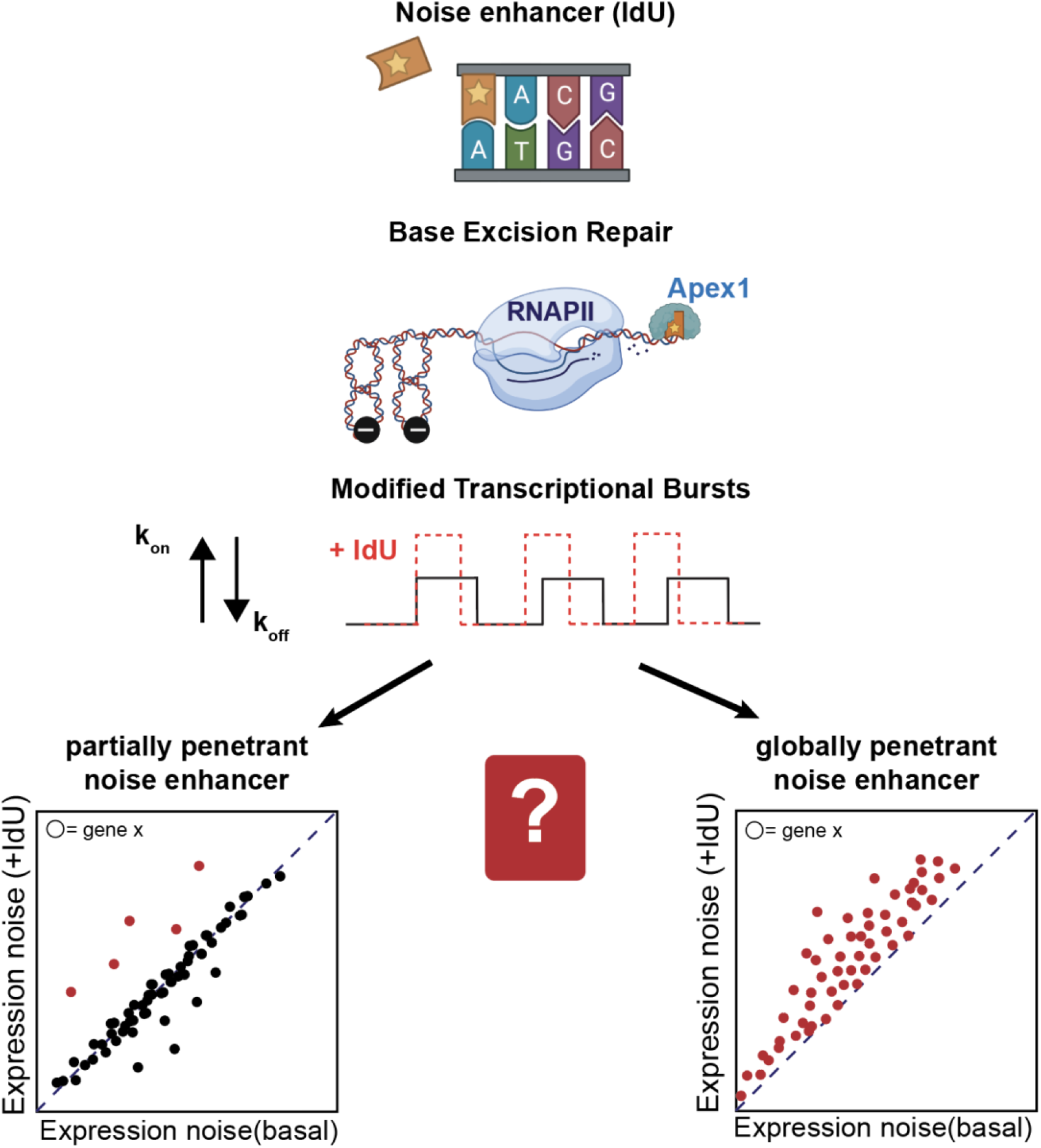

## INTRODUCTION

Cell-to-cell variability is an unavoidable consequence of the biochemical processes occurring in individual cells ^1^. While some portion of cell-to-cell variability arises from extrinsic factors (e.g., cell size, cycle phase, or microenvironment), a substantial body of literature has demonstrated that, in isogenic populations of cells—particularly mammalian cells—a large fraction of the variability originates from intrinsic sources, such as stochastic fluctuations (noise) in transcription ^2,3^. These intrinsic stochastic fluctuations can be quantitatively accounted for by the ‘toggling’ of genes between active and inactive expression periods leading to episodic bursts of transcription— commonly referred to as the two-state random-telegraph model of gene expression ^4–7^. The resulting transcriptional bursts are amplified upon nuclear export and cytoplasmic mRNA processing ^8^. Ultimately, these stochastic transcriptional fluctuations can generate substantial cellular variability, which has been implicated in cell-fate specification decisions ranging from HIV latency to cancer ^9–11^.

Although a few specific roles of transcriptional noise have been elucidated, it remains unclear how broadly this phenomenon impacts generalized physiological processes, particularly in eukaryotic and mammalian systems. Isolating the physiological roles of expression noise would typically require one to modulate noise independent of the mean level of expression ^12^. While this orthogonal perturbation of noise has been achieved in some notable cases ^13–15^, in practice, it is non-trivial to implement. The complication stems in part from the fundamental linkage between the variance (σ^2^) and the mean (μ) for most physical processes. For example, the common metric for quantifying expression noise, the coefficient of variation (CV), defined as σ/μ, typically scales inversely with mean, such that CV decreases as the mean increases ^16^. Consequently, the normalized variance, or Fano factor (σ^2^/μ) is often used to compare noise for processes with different mean values as it does not scale with the mean ^17,18^. Regardless of the metric, orthogonal noise modulation remains challenging and generalized approaches to modulate noise would be useful physiological probes.

To break the 1/μ dependence of CV and orthogonally modulate noise, specific autoregulatory architectures (e.g., feedback and feedforward) are typically required ^19,20^. However, small molecules called “noise enhancers” can also generate increased expression noise *without* altering the mean-expression level ^21^ by a process known as homeostatic noise amplification ^22^; a notable contrast to transcriptional activators, which increase mean-expression levels. We reported the molecular mechanism for one particular class of noise-enhancer molecule, pyrimidine-base analogs such as 5’-iodo-2’-deoxyuridine (IdU), and used scRNA-seq to show that IdU increase noise. However, scRNA-seq, despite its utility in measuring genome-wide expression ^23^, suffers from well-established issues of technical noise due to small inputs of RNA, amplification bias, and differences in capture ability ^24^ that could obscure the *penetrance* of IdU-mediated noise enhancement.

To better understand the genome-wide effect of IdU, here we reanalyzed scRNA-seq data using different normalization algorithms that minimize extrinsic and technical noise ^25^. Each algorithm identified a different proportion of the genes exhibiting amplified noise and—in the fraction of genes with amplified noise—substantial differences in the magnitude of noise amplification. To validate scRNA-seq measurements, we employed single molecule-RNA FISH (smFISH)—the gold-standard for mRNA quantification due to its high sensitivity for mRNA detection ^26^—to probe a panel of genes—from across the transcriptome—that displayed the greatest differences in noise between different scRNA-seq algorithms. Collectively, these analyses indicate that IdU is a globally penetrant noise-enhancer molecule and could, in principle, be a candidate to probe the physiological roles of expression noise for diverse genes of interest.

## RESULTS

### Alternate scRNA-seq normalization algorithms generate differing profiles of expression noise indicating noise amplification (ΔFano > 1) for ~90% of expressed genes

To examine how different scRNA-seq normalization methods influence the quantification of transcriptional noise, we employed six commonly-used normalization algorithms to analyze scRNA-seq data from IdU-treated versus DMSO-treated (control) mouse embryonic stem cells (mESCs) ^22^. In this dataset, cells were treated with the established noise-enhancer molecule IdU ^21,27^, noise was quantified by examining CV^2^ relative to the mean, and quality control was performed using Seurat ^28^, prior to all normalizations, to filter out low-expressing cells. Initially, we attempted to account for differences in sequencing depth between IdU and control samples using a rudimentary random subsampling (or downsampling) algorithm ^29^ to normalize control and IdU-treated samples to the equal read depths. The rationale for random subsampling is based upon reports that lower sequencing depth can be associated with increased variance ^30^, and subsampling reads to normalize the sequencing depth between samples can, in principle, correct for this. Surprisingly, downsampling suggested that IdU-mediated noise amplification was only partially penetrant, with ~78% of transcripts exhibiting amplified noise measured by Fano factor—this is compared to over 96% of transcripts exhibiting amplified noise when a standard log normalization without downsampling is used (**Fig. S1A-B**).

To determine if these subsampling results were consistent with other algorithms, we compared the results to five other established scRNA-seq normalization algorithms (**Fig 1**): *SCTransform* ^31^, *scran* ^32^, *Linnorm* ^33^, *BASiCS* ^34^, and *SCnorm* ^35^. *SCTransform* is a commonly used normalization algorithm that accounts for changes in depth and includes a variance-stabilization transformation step. *scran* and *BASiCS* are widely used normalization methods that use “size factors” to eliminate technical noise while maintaining biological heterogeneity. *Linnorm* utilizes homogenously expressed genes and scales the reads accordingly. *SCnorm*, similar to *SCTransform*, groups genes based on count-depth relationships and uses quantile regression to generate normalization factors.

**Figure 1:**
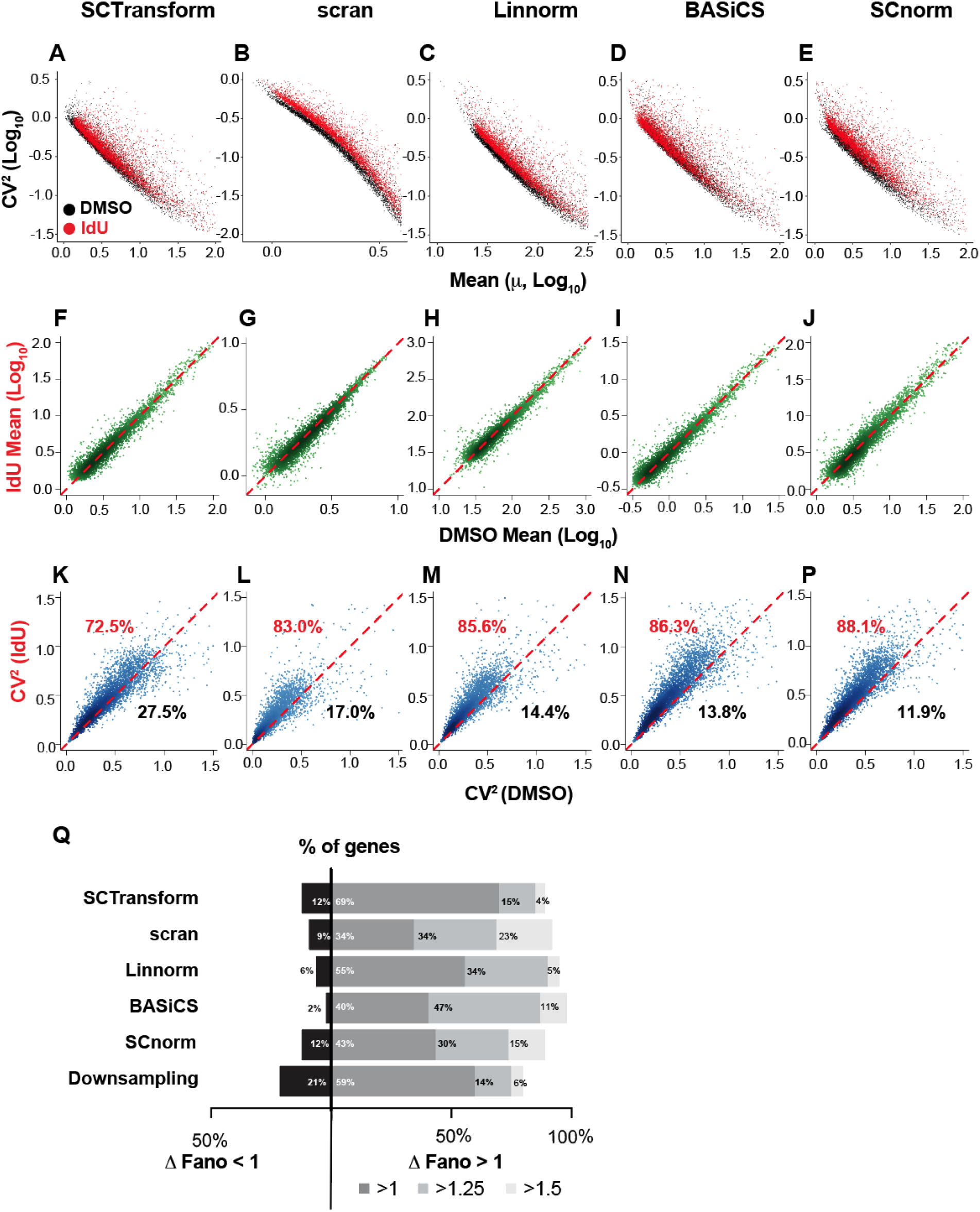
Common scRNA-seq normalization algorithms generate different quantifications of mRNA noise. scRNA-seq analysis of CV^2^-vs-mean for ~5,000 transcripts in mESCs treated with IdU (red) or DMSO control (black) as analyzed by commonly used normalization algorithms: (**A**) SCTransform, (**B)** scran, (**C)** Linnorm, (**D**) BASiCS, or (**E**) SCnorm. (**F-J**) Mean expression for each of the ~5,000 transcripts in presence and absence of IdU using each normalization algorithm; none of the normalizations algorithms generate substantial changes in mean expression for IdU-treated cells. (**K-P**) CV^2^ for each transcript in presence and absence of IdU using each normalization algorithm; different algorithms generate substantially different fractions of transcripts with amplified noise ranging from ~70% of transcripts with amplified noise (SCTransform) to ~88% of transcripts with amplified noise (SCnorm). (**Q**) Quantification showing percentages of transcripts with Fano factor fold change (Δ) <1, >1, >1.25 or >1.5 with IdU treatment. Calculations from all scRNA-seq normalization algorithm are shown (see Fig. S1 for Downsampling analysis).

Despite their substantially different technical schemes, analysis with each of these normalizations indicated that IdU induces a substantial amplification of noise (CV^2^) for most expressed genes (**Fig. 1A–E)**. The IdU-induced noise amplification appeared to be homeostatic (**Fig. 1F-J**), with mean-expression levels largely unchanged by IdU under all algorithms. However, each algorithm calculated a somewhat different percentage of expressed genes with increased CV^2^ ranging from 72% to 88% of genes exhibiting increased noise (**Fig. 1K–P**). Quantification of noise by analysis of the normalized variance (i.e., Fano factor), which in principle eliminates the scaling dependence on mean-expression level, showed a similar profile **(Fig. 1Q)**. Whilst every algorithm showed a majority of expressed genes exhibit amplified noise by Fano factor, each calculated a different penetrance (i.e., percentage of noise-amplified genes) as well as substantial differences in the magnitude of noise amplification among those transcripts with amplified noise.

Overall, scRNA-seq analysis by all normalization algorithms showed that IdU homeostatically amplifies noise for the majority of expressed genes without observable changes in the mean-expression level. However, each scRNA-seq normalization algorithm calculated a different transcriptional *penetrance* for IdU noise amplification, indicating that additional analysis of transcriptional noise via an independent (i.e., non-scRNA-seq approach) was warranted.

### RNA quantification by smFISH indicates widespread penetrance of IdU-induced noise amplification

To directly quantify RNA levels in individual cells using a non-sequencing-based approach, we employed smFISH, a well-established imaging method to assess noise and the gold standard for quantitative assessment of mRNA expression ^36^. While smFISH enables quantification of mRNA abundance in individual cells, it is a relatively low-throughput method requiring distinct fluorescent probes and image quantification for each transcript species of interest. Consequently, to generate a pseudo-global view of transcription using smFISH, we selected a panel of eight genes **(Fig. S1)** that satisfied two criteria: (i) they displayed the greatest difference in noise amplification as calculated by the different RNA-seq algorithms (**Fig. S1C-D**), and (ii) the genes were compatible with smFISH probe design (i.e., a minimum of 30 probes were predicted to hybridize to the transcript). The selected genes spanned a wide range of gene-expression levels (i.e., across 2-Logs and genome locations) (**Fig. S1E**) ^37^. We also included previously reported smFISH data ^22^ from a ninth gene, *Nanog*, to benchmark the smFISH analysis. smFISH probe sets for each gene in the panel were generated, cells were imaged in the presence/absence of IdU, and images (**Fig. 2A, S2A**) segmented and analyzed using FishQuant ^38^. Extrinsic noise was filtered by analyzing cell size (i.e., cell-area distribution in pixel number) by Kolmogorov-Smirnov test and excluding cells in the upper and lower tails of cell-size distribution (**Fig. S2B**). For each gene, the per-cell mRNA abundance was quantified for ~200 cells in order to generate the distribution of each mRNA per cell, in the presence and absence of IdU (**Fig. 2B**). This analysis revealed greater variance in per-cell mRNA levels in IdU-treated samples than controls, and the calculated CV^2^ values verified that noise increased for most genes (**Fig. 2C**). Direct comparison of mRNA CV^2^ values showed significant noise amplification for 8 of the 9 genes in the representative panel, without substantial changes in mean (**Fig. 2D, S3A**), in agreement with the scRNA-seq analysis. Notably, *Sox2* appeared to be an exception that exhibited a reduction in mean expression (see Discussion).

**Figure 2:**
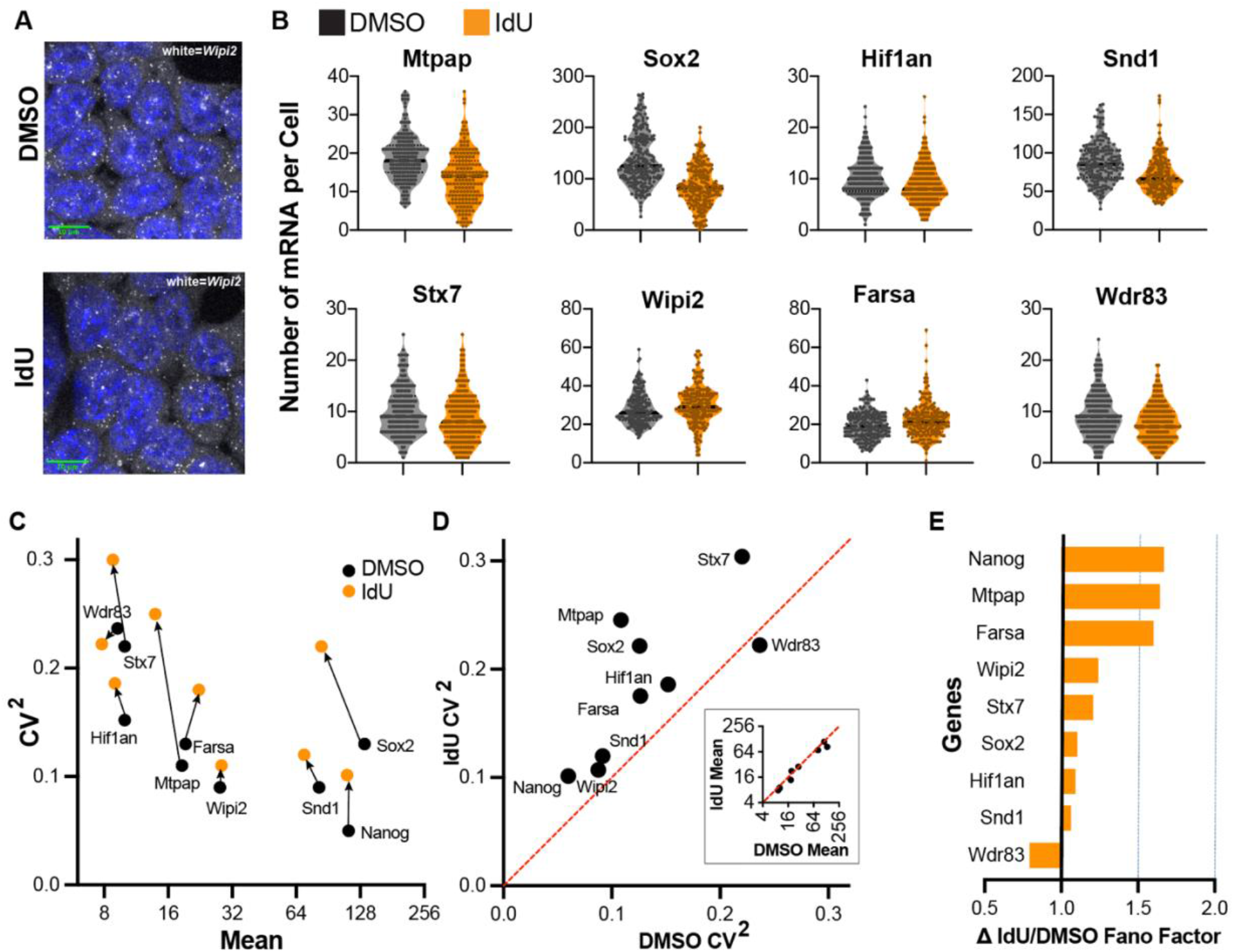
smFISH analysis of IdU-induced noise amplification for a subset of genes from accoss the transcriptome. (**A**) Representative smFISH images of *Wipi2* transcripts (white dots) in mESCs treated with DMSO (too) or IdU (bottom). Nuclei are stained with DAPI (blue). (**B**) Distribution of per-cell mRNA transcripts for eight representative genes +/-IdU. (**C**) CV^2^-vs.-mean analysis as calculated from smFISH +/- IdU for each of the genes in the representative panel. (**D**) Direct comparison of CV^2^ +/- IdU as calculated from smFISH data. **Inset**: direct camprrison of I dU-induced channel in mean mRNAfrom smFISH data. (**E**) Fold change in Fano Factor for IdU-treated samples compared to DMSO control from smFISH data.

To ensure that noise amplification, as reported by CV^2^, could not be explained by changes in mean expression, we also analyzed the Fano factor calculated from the smFISH data (**Fig. 2E, S3B**). Fano factor analysis verified that all but one gene in the panel *(Wdr83)* exhibited IdU-induced amplification of noise (ΔFano factor > 1). The greatest increased in noise was for *Nanog, Mtpap,* and *Farsa* (ΔFano factor > 1.5) whereas *Sox2, Snd1, Stx7, Hif1an,* and *Wipi2* exhibited smaller noise amplifications.

To further verify IdU-mediated noise amplification, we tested if the mechanistic underpinnings ^21^ of homeostatic noise amplification were satisfied. Theory predicts that homeostatic noise amplification requires reciprocal changes in transcriptional bursting parameters ^21^; i.e., a decrease in transcriptional burst frequency with a corresponding but reciprocal increase in transcriptional burst size. Transcriptional burst frequency can be calculated by scoring the percentage of cells with active transcriptional centers (TCs) in smFISH images, and the burst size can be similarly calculated if other parameters, such as the decay rate of the specific mRNA, are also known ^39^. Consequently, we calculated burst frequency and burst size for two genes, *Sox2* and *Mtpap*, for which mRNA decay rates in mouse embryonic stem cells were known ^40^. In agreement with the model of homeostatic transcriptional noise amplification, IdU substantially decreased the burst frequency (**Fig. S4A**) and correspondingly increased the burst size (**Fig. S4B**) for both genes.

Overall, these smFISH data indicate that IdU generates a homeostatic amplification of transcriptional noise for 8 out of 9 genes in the representative panel taken from diverse expression profiles and locations in the genome.

## DISCUSSION

This study set out to determine if the pyrimidine base analog IdU, which is incorporated into cellular DNA and removed by the cellular base-excision repair (BER) surveillance machinery ^41^, is a globally penetrant noise-enhancer molecule. Analysis of scRNA-seq data using six different normalization algorithms indicated that the amplification of noise by IdU is globally penetrant and likely not an artifact of a particular scRNA-seq analysis algorithm. However, the scRNA-seq analysis also indicated that scRNA-seq algorithms generate variable genome-wide noise profiles (**Fig. 1**). To validate IdU’s genome-wide effect, we curated a panel of genes that exhibited high sensitivity to individual scRNA-seq normalization algorithms and represented both high and low expressing genes for downstream orthogonal analysis of transcriptional noise by smFISH. This approach revealed that IdU increased transcriptional noise for 8 out of 9 genes (**Fig. 2**) and that most published scRNA-seq algorithms (with the exception of rudimentary downsampling) were fairly accurate compared to the smFISH direct measurement (compare **Fig. S1C** to **Fig. 2E**).

One technical limitation of this study is that smFISH analysis is necessarily low throughput and limited to the subset of genes for which good probe sets can be designed, which limits the spectrum of measurements and the number of genes that can be analyzed. We attempted to mitigate this by exploring genes from across the expression spectrum (**Fig. 2C**), and by analyzing a substantial number of cells per treatment (>170 cells per treatment **after** filtering extrinsic noise). Thus, our data indicate that IdU likely acts as a noise-enhancer molecule for a large fraction of genes irrespective of their mean-expression level.

Notably, our smFISH analysis did not detect an increase in noise for *Wdr83.* Considering that IdU increases noise via BER, a genome-wide surveillance pathway, it may be surprising that any gene fails to exhibit an increase in noise. It is possible that the low-throughput nature of smFISH may have obscured IdU-induced noise amplification. However, a potential explanation for the selective response lies in the putative mechanism by which BER increases noise via DNA topology changes. Specifically, the BER enzyme AP endonuclease 1 (Apex1) generates DNA supercoiling which leads to an accumulation of RNA Polymerase II; when released, this amplifies transcriptional burst size; indeed, the IdU noise effect can be phenocopied by topoisomerase inhibitors, which increase DNA supercoiling ^22^. Consequently, regions which naturally have increased supercoiling and corresponding high levels of topoisomerase may be less sensitive to changes in topology caused by IdU and BER-induced supercoiling. It has been previously reported that Topologically Associated Domain (TAD) boundaries are regions that exhibit substantial supercoiling and are enriched in insulator binding protein CTCF ^42,43^, and that Toposimerase IIB is prevalent at CTCF sites ^44^. Intriguingly, *Wdr83* has two relatively unique features among the panel of genes tested: (i) it contains a relatively large CTCF binding region, and (ii) it is found at a TAD boundary ^42^.

Together these two features may be consistent with IdU acting as a topology-dependent global noise-enhancer molecule, as shown by scRNA-seq **(Fig. 1)** and smFISH **(Fig. 2)** analysis.

A second point of interest is the case of *Sox2.* Whilst most genes analyzed exhibited a homeostatic increase in noise—without a substantial change in mean—smFISH revealed that IdU induced a decrease in mean *Sox2* mRNA, though we note the change in mRNA numbers was less than two-fold. Notably, IdU did not alter single-cell *Sox2* protein levels in our previous analysis ^22^. It is possible that the reported negative-feedback regulation of *Sox2* ^45^ acts to buffer changes at the protein level.

From the practical perspective of quantifying expression noise, this study reveals that common analyses can fail to resolve quantitative changes in intrinsic expression noise, particularly for individual genes. A number of approaches have been proposed to overcome these types of scRNA-seq limitations, including elegant solutions using mathematical modeling methods to address technical variability without compromising quantification of biological noise ^46^, though these can be cumbersome to implement. The analyses herein indicate that it may be advisable to combine high-throughput analyses (e.g., scRNA-seq) with lower-throughput direct quantification (e.g., smFISH) to quantify changes in transcriptional noise for individual genes as the “ground-truth” may lie somewhere in between the results of each analysis. Regardless, both the scRNA-seq and smFISH data argue that IdU appears to be a global noise enhancer which could be leveraged to modulate noise without altering mean expression for the majority of genes.

## Acknowledgements

We thank L. Miranda for administrative support; R. Desai, G. Vasen, K. Delucas, and D. Lewis for technical guidance. Harrington and S. Kim for imaging guidance; K. Claiborn for editing and O. Rechavi for subsampling guidance. Data for this study were acquired at the Nikon Imaging Center at UCSF using NIH S10 Shared Instrumentation grant (1S10OD017993-01A1).

## Author contributions

GC, XC, and LSW conceived and designed the study. GC and XC designed the smFISH experiments. GC performed, analyzed, and curated the data. GC and BZ performed single-cell Seq normalizations and GC curated the data. LSW provided reagents and resources. GC and LSW wrote the paper.

## SUPPORTING FIGURES

**Figure S1:**
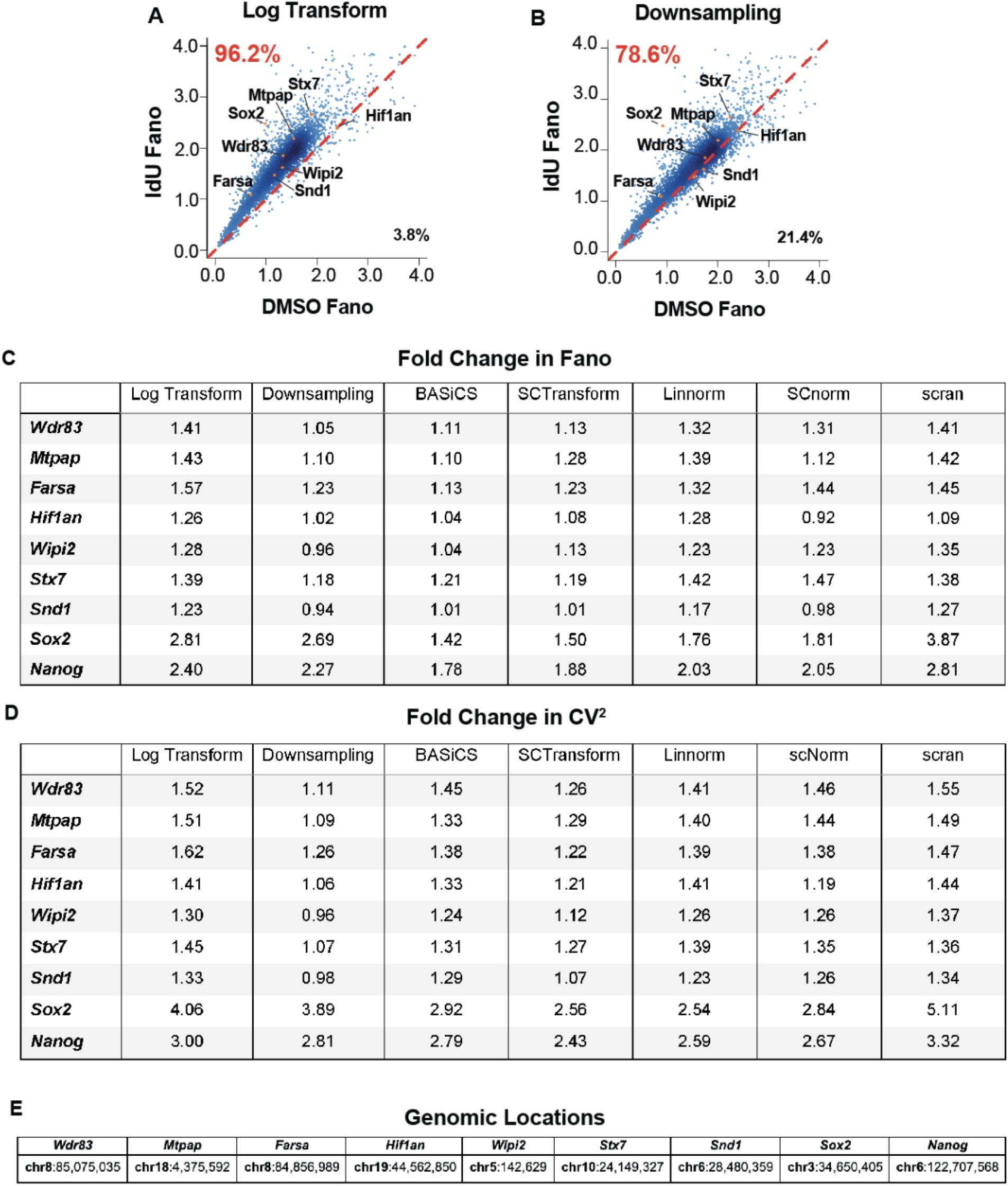
Single Cell RNA Sequencing algorithms show varying penetrance of IdU-induced amplification of expression noise. Related to Figure 1. Direct comparison of Fano value for different single cell sequencing normalizations. The 8 genes chosen for smFISH are labeled in **(A)** Data used from Desai et al., 2021. **(B)** Downsampling algorithm that matches DMSO sequencing depth with IdU sequencing depth using random subsampling. Table of values showing the fold change of IdU vs control for Fano (**C**) and CV^2^ (**D)** for the 8 genes selected for smFISH (together with *Nanog* from Desai et al. 2021) across the seven normalization algorithms. **(E)** Genomic locations of genes selected for smFISH.

**Figure S2:**
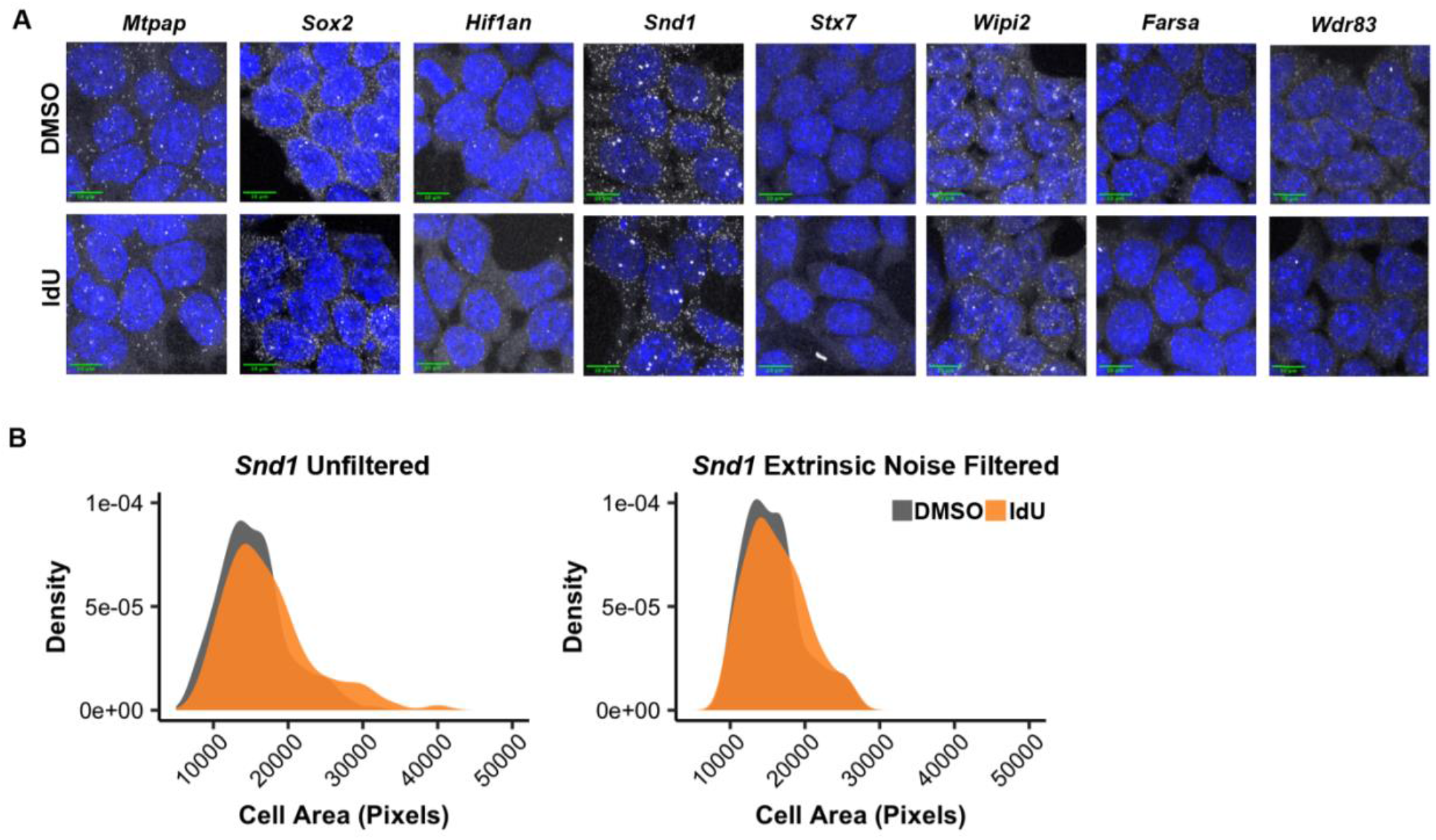
Representative images and extrinsic-noise filtering (by cell-size) for smRNA FISH. Related to Figure 2. (**A**) Representative smFISH images of mESCs treated with DMSO (top) or IdU (bottom) for 24 hours with DAPI stain (blue) and mRNA transcripts (gray) fluorescent labeled with TAMRA probe-set. Max intensity projections are shown. (**B**) Left: The cell size distribution of 194 DMSO treated cells and 196 IdU treated cells are significantly different (KS test p = 0.007). Right: the 5th and 95th percentile of the whole population were removed. The cell size distribution for the remaining 178 DMSO Treated cells and 174 IdU treated cells are not significantly different (KS test p=0.21)

**Figure S3:**
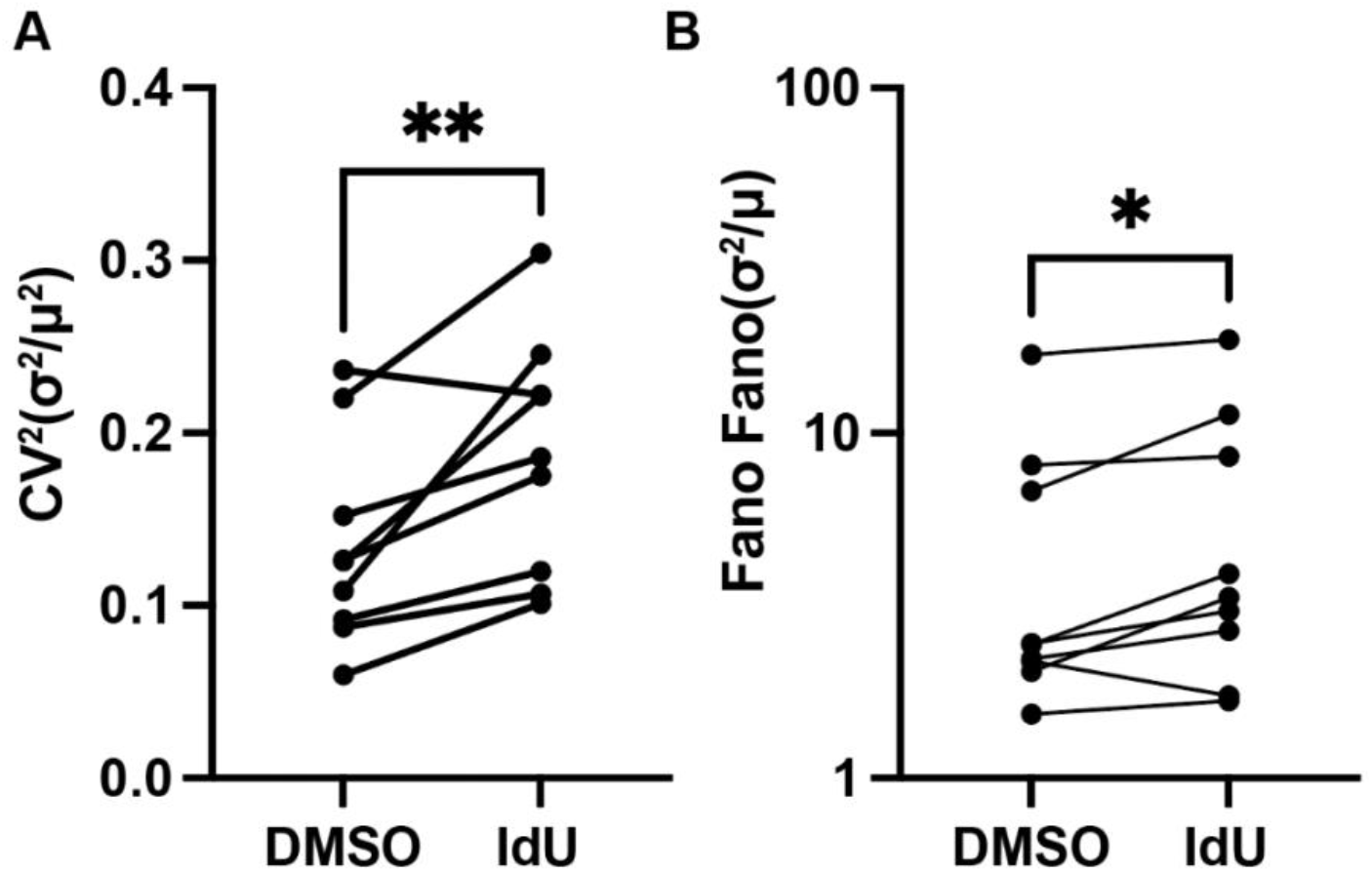
smRNA FISH shows significantly increased Fano Factor and CV^2^ with IdU Treatment. **(A)** Fano Factor comparison of samples +/- IdU compared using Wilcoxon matched-pairs signed rank test p =0.0117. **(B)** CV^2^ comparison of samples +/- IdU compared using Wilcoxon matched-pairs signed rank test p = .0078

**Figure S4:**
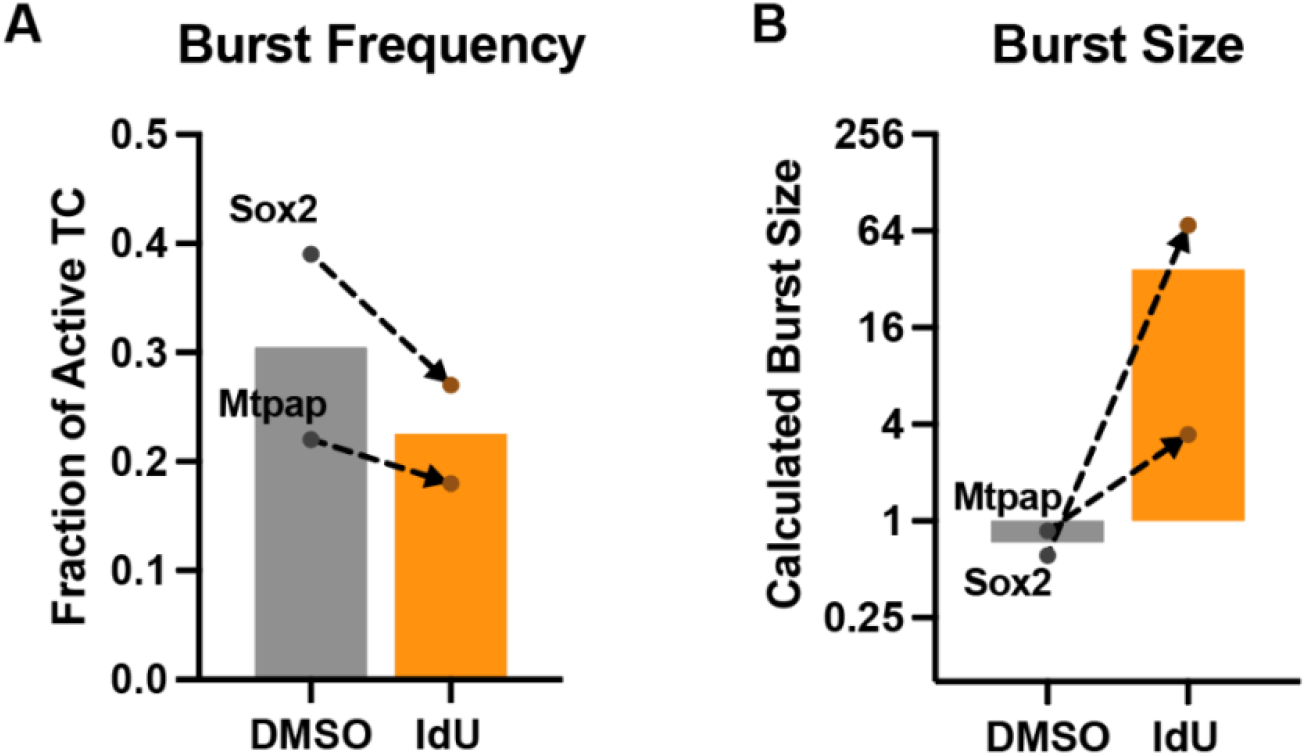
Changes in burst frequency and burst size with IdU treatment. **(A)** Transcriptional burst frequency for *Mtpap* and *Sox2* as calculated from the fraction of active transcription centers (TC) in smFISH images. **(B)** Calculated transcriptional burst size for *Mtpap* and *Sox2* (see Methods).

## Captions for supplementary table S1

**Table S1 (**attached separately)

Sequences of smRNA-FISH oligonucleotide probe for eight genes

## Data Availability

This study includes no new data deposited in external repositories.

## Star Methods

### Single-Cell RNA Sequencing Analysis

The scRNA-seq count matrix available at GSE176044_mesc_bulk_rnaseq_gene_counts was used (Data ref: Desai *et al,* 2021). Before applying each normalization method, both DMSO and IdU datasets were subjected to a quality control process. First, Seurat ^28^ was used to filter for high-quality cells using a minimum of 4000 detected genes, 10000 UMI counts, and <10% reads mapping to mitochondrial genes per cell.. The resulting count matrix was then further filtered by the BASiCS_Filter function from the BASiCS ^34^ R package with default parameters, which limited the analysis to genes with sufficient sequencing coverage for reliable noise quantification. The output count matrix consisted of 811 cells for DMSO and 732 cells for IdU across 4456 genes. This output was then used to run 5 normalization pipelines according to their protocols: *SCTransform* ^31^, *scran* ^32^, *Linnorm* ^33^, *BASiCS* ^34^, and *SCnorm* ^35^. BASiCS was run with default parameters and recommended settings (N=20000, Thin=20 and Burn=10000) using the horizontal integration strategy (no-spikes). The other packages were run with default parameters. To perform rudimentary downsampling **(Fig. S1B**) the python random.seed function was used to simulate an identical sequencing depth for both DMSO and IdU. 81% of DMSO reads were sampled, and 100% of IdU reads were subsampled. From this subsampled matrix, the same 4445 genes and 1543 cells included previously were used for further quantification steps.

### Single Molecule RNA FISH

Probes for the detection of transcripts were developed using the designer tool from Stellaris (LGC Biosearch Technologies) (Table S1) setting the minimum number of probes to 30 (TAMRA conjugated) for gene transcripts. 1×10^5^ mouse embryonic stem cells were seeded into each well of a gelatin-coated, 8-well Ibidi dish (cat: 80826) in 2i/LIF media. 24 hours following seeding, media was replaced with 2i/LIF containing 10mM IdU or equivalent volume DMSO. After 24 hours of treatment, cells were then fixed with DPBS in 4% paraformaldehyde for 10 minutes. Fixed cells were washed with DPBS and stored in 70% EtOH at 4°C for one hour to permeabilize the cell membranes. Probes were diluted 200-fold and allowed to hybridize at 37°C overnight. Wash steps and DAPI (Thermo, cat: D1306) staining were performed as described (https://www.biosearchtech.com/support/resources/stellaris-protocols). To minimize photo-bleaching, cells were imaged in a buffer containing 50% glycerol (Thermo, cat: 17904), 75 mg/mL glucose oxidase (Sigma Aldrich, cat: G7141), 520 mg/mL catalase (Sigma Aldrich, cat: C3515), and 0.5 mg/mL Trolox (Sigma Aldrich, cat: 238813). Images were collected on an inverted Nikon TiE microscope (Nikon) run using Micromanager 2.0 ^47^ equipped with a CSU-W1 Spinning Disk with Borealis Upgrade (Yokogawa, Andor), ILE Laser launch with 4 laser lines (450/488/561/646nm, Andor), quad-band dichroic ZT405/488/561/647 (Chroma), emission filters for DAPI (ET447/60), GFP (ET525/50), RFP (ET607/36), and Cy5 (ET685/40) (Chroma), piezo XYZ stage (ASI), and Zyla 4.2 CMOS camera (Andor), using a Plan Apo VC 60x/1.4 Oil objective (Nikon). Approximately 10 XY regions of interest were randomly selected for each condition. For each image, XY pixel size was 108nm/px, and a Z-step size of 250nm was used with over 60 image planes to fully cover the tissue. Image analysis and spot counting was performed using FISH-quant ^38^. Cells were manually segmented and analysis was conducted on cells of a similar size to minimize extrinsic noise. Active transcription sites were measured according to ^39^. Burst size was calculated as previously reported ^8^ using the following equation: Burst Size = (λ_mRNA_* μ_mRNA_)/k_off_ where k_off_ = 1- (% of active TC).

